# Contrasting patterns of foraging behavior in Neotropical stingless bees using pollen and honey metabarcoding

**DOI:** 10.1101/2023.06.06.543880

**Authors:** Aline C. Martins, Carolyn E. B. Proença, Thais N. C. Vasconcelos, Antonio J. C. Aguiar, Hannah C. Farinasso, Aluisio T. F. de Lima, Jair E. Q. Faria, Krissya Norrana, Marcella B. R. Costa, Matheus M. Carvalho, Rodrigo L. Dias, Mercedes M. C. Bustamante, Fernanda A. Carvalho, Alexander Keller

## Abstract

Stingless bees are major flower visitors in the tropics, but their foraging preferences and behavior are still poorly understood. Studying stingless bee interactions with angiosperms is methodologically challenging due to the high tropical plant diversity and inaccessibility of upper canopy flowers in forested habitats. Pollen DNA metabarcoding offers an opportunity of assessing floral visitation efficiently and was applied here to understand stingless bee floral resources spectra and foraging behavior. We analyzed pollen and honey of three distantly related species of stingless bees, with different body size and social behavior: *Melipona rufiventris, Scaptotrigona postica* and *Tetragonisca angustula*. Simultaneously, we evaluate the local floristic components through seventeen rapid botanical surveys conducted at different distances from the nests. We discovered a broad set of explored floral sources, with 46.3 plant species per bee species in honey samples and 53.67 in pollen samples. Plant families Myrtaceae, Asteraceae, Euphorbiaceae, Melastomataceae and Malpighiaceae dominated the records, indicating stingless bee preferences for abundant resources that flowers of these families provide in the region. Results also reinforce the preference of stingless bees for forest trees, even if only available at long distances. Our high-resolution results encourage future bee-plant studies using pollen and honey metabarcoding in hyper diverse tropical environments.

## Introduction

Plant-pollinator interactions mediate most flowering plant reproduction, maintaining terrestrial ecosystems and crops^1^. The current decline in pollinator abundance and diversity worldwide threatens pollination services, with direct consequences to nature conservation and food security^2^. Therefore, understanding the interaction between pollinators and flowering plants became crucial, since it can provide an information framework to subsidize conservation policies and decisions in a changing world. The most important group of animal pollinators – the bees – totally depend on floral resources to complete their life cycles^3^, while 87.5% of the animal-pollinated flowering plants depend on bees to reproduce^4^. Interaction between bees and angiosperms has been a major research focus in the last two decades, due to the massive effects of insect and bee declines^5^ threatening ecosystem services and food security^6,7^. In this context, pollen, and honey DNA metabarcoding emerged as an efficient technique to identify plant taxa visited by pollinators based on samples extracted from bees’ bodies or nests^8–11^. DNA metabarcoding of pollen and honey has been largely applied to temperate systems, and recently to (sub) tropical species of stingless bees^12,13^.

The stingless bees (Apidae, tribe Meliponini) comprise c. 500 species of eusocial bees, most of which occur in tropical America (more than 400 species), but also in Africa, Asia and Australia^3,14^. The role of stingless bees as pollinators of the neotropical flora could become even more relevant under climate change scenarios, if the predicted expansion of warmer temperatures pushes the distribution of predominantly temperate bees, such as *Apis* and *Bombus*, into cooler regions^6,15^. Although domestication is still restricted to a few species, stingless bees are also explored commercially for honey production ^16^, which can reduce the use of introduced honey-bees and their impact on native species in these regions^12^

As all social bees, stingless bees show a predominantly generalist pattern of floral exploitation, i.e. they visit a large number of species in several plant families, supposedly disregarding specific floral traits^17^. Particularities exist though, since stingless bees exhibits a huge diversity of body size (1.8 to 13.5 mm)^3^ flight distance (0.3 to 3 km)^18^ and foraging behavior^19^. However, the breadth of the stingless bees diet – that is, how many different species of flowering plant they forage on – and the extent of their role as pollinators of tropical plants - are still largely open questions. Most studies aiming to answer some of these questions faced some methodological difficulties due to inaccessibility of visited flowers, which mostly occupy upper canopy strata, especially in rainforests^20^, and hyper diversity of tropical plants, thus hampering easy identification through bee-pollen morphology^21,22^.

In this study, we explored the diet breadth of three distantly related species of stingless bees of different body sizes and flight ranges in a hyper-diverse tropical ecosystem, the Cerrado savannas of central South America. The Cerrado is the most species-rich savanna in the world and a hotspot of biodiversity^23^; the flora encompasses >13.000 native species^24^ a highly patchy vegetation with several different physiognomies, ranging from grasslands, marshlands, and typical savanna to closed canopy riverine forests along waterways^25^. The proportion of pollinator-dependent species in the Cerrado flora is still unknown, although this number is likely to be similar to that of tropical forests^4^, with some authors estimating c. 60% of angiosperm species being bee dependent^26^.

We analyzed pollen and honey from the pots of the nests (henceforward pot-pollen and pot-honey respectively) from three commonly managed stingless bee species (*Melipona rufiventris, Scaptotrigona postica* and *Tetragonisca angustula*) native to the Cerrado to investigate: (i) How broad is the floral resource exploitation of stingless bees in a hyper-diverse flora? (ii) Which plant species and families are the most important sources of pollen and/or nectar for stingless bees in the area? (iii) What can pollen and honey metabarcoding reveal, when combined with floristic surveys of the area, about stingless bees foraging behavior, particularly foraging distances, and floral preferences? We also discuss how efficiently pollen and honey metabarcoding identified plants visited by bees in the area, considering the low DNA sequence coverage of neotropical plant species in public databases^27^, and the potential role of this technique in improving ecological understanding of bee-plant interactions in the tropics.

## Material and Methods

### Study site

Our study was conducted in the Ecological Reserve of the Brazilian Institute of Geography and Statistics (IBGE) (15°56′41″ S and 47°53′07″ W) that, together with the contiguous Brasilia Botanic Garden and the University of Brasília Experimental Field Station, preserves an area of c. 10,000 ha of native Cerrado in the Distrito Federal, Brazil. The IBGE reserve was chosen as a study site for being one of the most well-studied areas of Cerrado, with good prospects of building a relatively robust plant DNA reference library, a requirement for our analyses (see below); it occupies a central position within the Cerrado Biome. The climate in the area is typical tropical savanna climate (Aw Köppen classification system) with dry winters and rainy summers with an average annual precipitation of 1453 mm, altitude ranges from 1048 to 1160 m. The IBGE reserve contains the main vegetation types typical of the Cerrado domain: savannas (*cerrado* sensu stricto), palm swamps (*veredas*), grasslands (*campo limpo* and *campo sujo*) and riverine forests (*mata de galeria*), surrounded by natural and agricultural areas (Figure 1). This habitat heterogeneity results in high plant biodiversity. The last published floristic survey in the area recorded 1798 species of angiosperms, of which 1457 are native, distributed in 138 families and 724 genera^28^.

**Figure 1.**
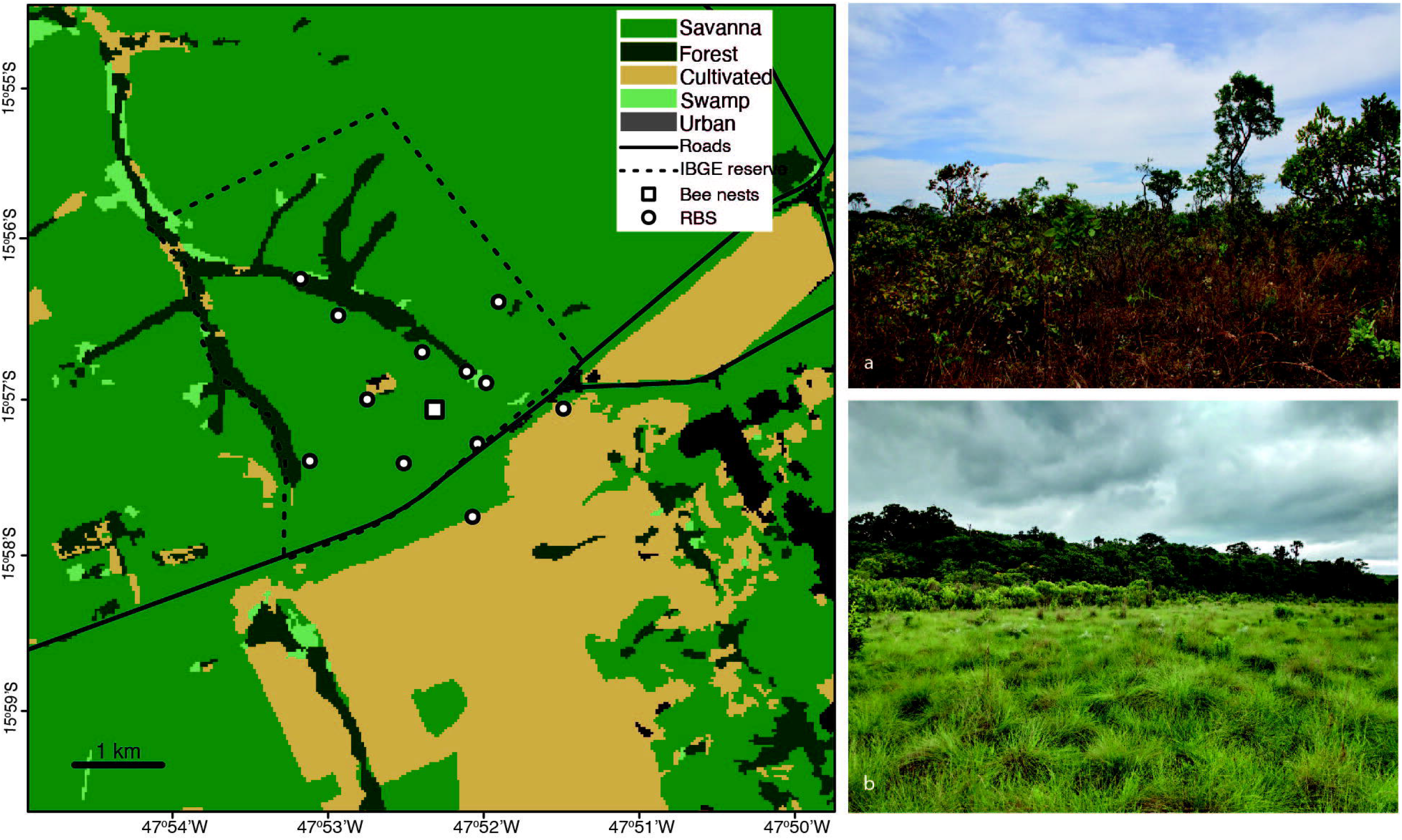
Map of the IBGE reserve and surroundings showing the location where bee nests were installed and the locations of Rapid Botanical Surveys. The image also shows main vegetational types, i.e. cerrado savanna, riverine forests, swamps, cultivated and urban areas. Photographs depict a. cerrado savanna vegetation type (Photo author: ACM) and b. area of transition between grassland and riverine forest (Photo author: AJCA). Vegetation cover: MapBiomas (www.mapbiomas.org). Reserve delimitation: IBGE.

### Stingless bee species and nest material sampling

Three native species of stingless bees were chosen for our study: *Melipona rufiventris, Scaptotrigona postica* and *Tetragonisca angustula*. Choice of species was guided by three key aspects: body size, differences in foraging behavior, and phylogenetic relationships. Melipona rufiventris is the largest with a body length of c. 9.5 mm, *S. postica* has an intermediate body size varying from 5.7 to 6 mm, and *T. angustula* is amongst the smallest stingless bees with a total body length of c. 4 mm. Phylogenetically, the three genera are not closely related, i.e. they are not sister groups^29^. *Melipona rufiventris* is typically found in the more open vegetation types of eastern and central Brazil, *Scaptotrigona postica* occours in a broader region in Central, Northeast and southeast Brazil, also associated with open vegetation, while *Tetragonisca angustula* is widespread in the Neotropics (Mexico to South America)^14^.

The three species are commonly managed by local beekeepers and were chosen also for their relatively easy management in artificial colonies (Figure S1). The decision to use artificial colonies for sampling in our study relied on four main points: 1. to preserve the natural bee community in the area, not destroying any nests for sampling; 2. to facilitate sampling, as pollen and honey are stored in accessible compartments in the wooden box; 3. to facilitate the access to the colonies, which in natural conditions would be randomly distributed, depending on availability of cavities, and 4. to make sure nests would have a strong population and enough pot-pollen and pot-honey for sampling. Eighteen pre-established nests were installed: three of *M. rufiventris*, eight of *S. postica* and seven of *T. angustula*. The nests were installed at a distance of about 5 m from each other and c. 150 cm above ground level, in typical savanna or *cerrado sensu stricto*^28^ where most species are subshrubs, shrubs or small trees.

Nests were moved to the study area eight weeks prior to the first sampling to allow bees time to start accumulating pollen and honey from local species in the artificial nests. Pot-pollen and pot-honey samples were collected from the nests (Figure S3) once every 15 days for five months (July 2019 – November 2019). This period started at the height of the dry season, moved through the transition between dry and wet seasons and ended at the beginning of the wet season. Samples were always collected from new pots – that is, those built in between two subsequent sampling events. Micropipettes (1000 uL) were used to collect honey from the pots, while pollen was collected with plastic straws, which perforates the pollen mass while collecting it at the same time. Samples were subsequently stored in falcon tubes and stored in a -20° C freezer until extraction. In total, 191 samples (115 of pollen and 75 of honey) were collected from the three species: 29 of *M. rufiventris*, 81 of *S. postica* and 74 of *T. angustula*

### Metabarcoding protocol

Extractions of DNA of pollen samples from pot-pollen and pot-honey follow different methodologies, due to the different natures of the samples. For pot-honey, we extracted DNA using the Machery-Nagel (Düren, Germany) NucleoSpin Food Kit; for pot-pollen we used the Machery-Nagel (Düren, Germany) NucleoSpin Plant II.

#### Pot-Pollen DNA extraction

To extract pollen genomic DNA, we added to the pooled samples (weight ranging from 0.1 g to 2 g) 4 mL of deionized and autoclaved water, and homogenized it using a vortex. We then placed 200 μL of this emulsion in a 1.5mL microcentrifuge tube, and centrifuged it for 15 minutes at 8000 rpm. We discarded the supernatant material, froze the pellet obtained in liquid nitrogen, and then used mortar and pestle to break the pollen exine and the NucleoSpin Plant II Kit to promote cell lysis and to isolate the DNA according to the manufacturer’s instructions.

#### Pot-Honey DNA extraction

To extract pollen genomic DNA from honey, we added deionized and autoclaved water to the samples until the volume of each sample tube reached 1.5 mL. We incubated the tubes at 65 □ °C for 30□ min and, over that period, inverted the tubes slowly to homogenize the material. We then pooled the honey samples collected from the same nest and the same day by pouring them into falcon tubes, to which deionized and autoclaved water was added until completing 10 mL. Afterwards, we centrifuged these pooled samples for 15 min at 5000 rpm and discarded the supernatant material. Each precipitated pooled honey sample was resuspended in 200 μL deionized and autoclaved water and placed in a 1.5-mL microcentrifuge tube. This procedure was done twice. Finally, we centrifuged the samples for 15 min at 5000 rpm, discarded the supernatant material, dried the pellet in a drying cabinet at 35°C, and then ground the samples inside the microcentrifuge tube using micro-pestles and liquid nitrogen. We then used the NucleoSpin Food Kit to promote cell lysis and to isolate the DNA according to the manufacturer’s instructions.

The protocol of amplification utilizes a dual-indexing strategy^9^ to amplify the ITS2 region, using the primers ITS-S2F and ITS4R. Primer sequences, references and other amplification methodological details can be found in^9^ and^30^. The triplicate PCR reactions were combined per samples, well mixed and checked on 1% agarose gel using 5 uL of the combined products for quality. PCR products of each sample were normalized to ensure more equalized library sizes using the SequalPrep Normalisation kit (Invitrogen, CA, USA) according to the manufacturer’s protocol. The multiplex-index samples were pooled and then submitted to quality control and quantification to ensure the correct fragment size has been amplified with a Bioanalyzer High Sensitivity DNA Chip (Agilent Technologies, CA, USA) and a dsDNA High Sensitivity Assay on the Qubit Fluorometer. For library dilution, we followed the Illumina Sample Preparation Guide for a 2 nM library and a 5% PhiX control was added in order to increase quality. In addition, the reagent cassette of the sequencing kit was spiked with the Read1, Read 2 and index primers according to Sickel et al. (2015). Sequencing was then performed on the Illumina MiSeq system at the University of Würzburg. Sequence data are available at NCBI (Bioproject 976708).

### Bioinformatic data analyses

We used VSEARCH v2.14.2^31^ to join paired ends of forward and reverse reads and to remove reads shorter than 150bp, quality filtering (EE < 1)^32^, *de-novo* chimera filtering (following UCHIME3)^33^, and determination of amplicon sequence variants (ASVs)^33^, as previously done for pollen metabarcoding networks^12^. Reads were first directly mapped iteratively with global alignments using VSEARCH against several floral ITS2 reference databases for the study region and an identity cut-off threshold of 97%. A reference library of ITS2 sequences of all plant species recorded from IBGE was built from sequences available on GenBank. This primary database was then curated to remove voucherless entries for greater trustworthiness. Remaining unclassified sequences were then tracked by iterative searches against geographically broadening public sequence reference data, i.e., species lists of the flora of the Distrito Federal, then the large, neighboring state of Goiás, and lastly the entire Cerrado biome flora to increase completeness of reads. These reference databases were created with the BCdatabaser^34^ from GenBank entries given above mentioned species lists and default parameters (length between 200 and 2000 bp, maximum nine sequences per species). For still unclassified reads, we used SINTAX^35^ to assign taxonomic levels as deep as possible using a global reference database^36^. After classification, we performed plausibility checks according to geolocation and phenology with the results to verify validity. Thirteen species were automatically matched to genus level only but were attributed to species based on being the only species of the genus to occur in the Distrito Federal.

### Floristic surveys and vegetation characterization

To improve our knowledge of the flora surrounding the nests, we conducted Rapid Botanical Surveys (RBS) in small plots that were demarcated *in loco* as homogeneous to vegetation type. These plots were exhaustively surveyed for all flowering plant species of all life forms, fertile or not, by a team of 3-5 researchers, where one was the booker, i.e. the most experienced person in the group, who identified the plants in the field and discarded duplicated species; other team member collected and pressed the vouchers (for additional methodological details see^37^.

Eleven RBS plots had been initially chosen to correspond to one plot near the nests (henceforward nest plot) and ten other plots established at the vertices of two pentagons; the inner pentagon was established with its vertices at 700m from the nests and the outer pentagon with vertices at 1500m from the nests. These distances were chosen based on the literature of the flight capabilities of other stingless bees^18^. These eleven RBS plots mostly fell in areas of well-preserved savanna within the IBGE Reserve, ranging from the more open, grass and herb-rich areas with few shrubs and trees (*campo sujo*), to dense savanna woodland (*cerradão*); one outer pentagon plot fell in disturbed cerrado and another in heavily degraded secondary vegetation out of the IBGE. Because none of the plots fell in riverine gallery forest, we included six additional RBS plots in this vegetation type: three in the riverine gallery forest nearest to the nests (Nascente do Roncador, c. 630m from the nests), and three in a more distant gallery forest (Ponte do Corujão, c. 2070m from the nests), measured as the crow flies, thus a total of 17 RBS plots. Lastly, we also surveyed the plants and weeds growing in the ornamental gardens associated with the Main Building and Seat of the Reserva Ecológica do IBGE, which is located c. 650m from the nests. All specimens collected in RBS inventories were deposited in the UB Herbarium (University of Brasilia) and the records are available online in the Species Link Network (https://specieslink.net/search/) by searching on the collector name “Projeto Barcode Cerrado”.

### Data Integration

The 30 most abundant plant species in the pollen and honey samples were classified by ubiquity (i.e., presence in pollen or honey samples of two or all bee species). We then crossed this information with data from the RBS floristic surveys: distance from the nests: i.e., if they were sampled at nest plot, inner pentagon plots, outer pentagon plots, nearest or furthest gallery forest plots, or the gardens. These 30 species were also characterized from the literature in terms of their offered resources (e.g. pollen, nectar, oil, resin), their habitat (savanna, forest or cultivated/weedy) and habit (trees, shrubs, subshrubs, hemiparasites) (Table 1).

**Table 1.**
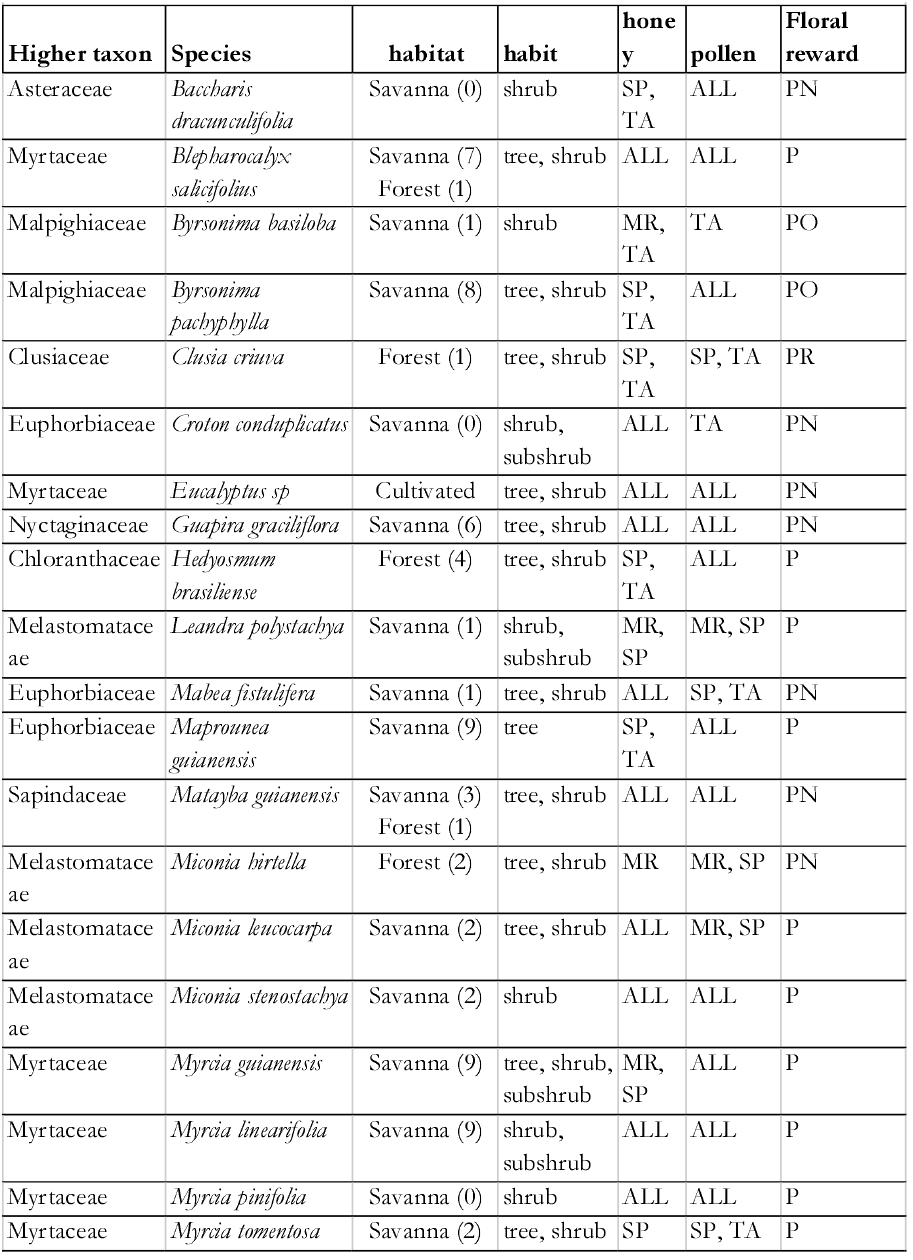
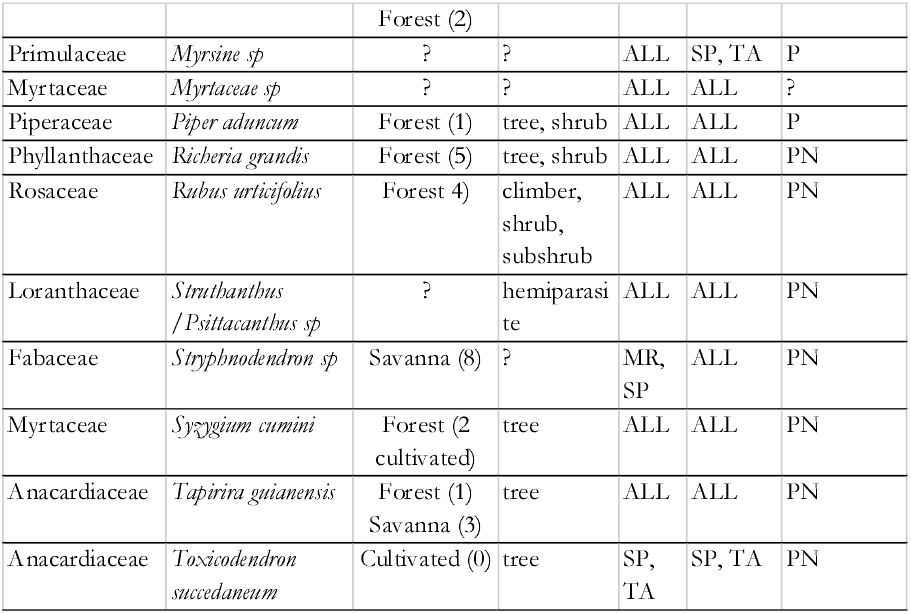
Thirty most frequent taxa in ASVs, their habitats and habits (tree, shrub, subshrub, climber, hemiparasite), presence in pollen or honey and floral resource offered (P: pollen; N: nectar; O: oil; R: resin). Habitat data from floristic inventory in this study; numbers in parenthesis represent a record in each RBS plot (forest plots surveyed: 6; savanna plots surveyed: 11). Habit data from Flora & Funga do Brasil (2023).

### Statistical analysis of pollen and honey samples

Data was processed for analyses using R 4.2.2^38^ and the packages phyloseq^39^ vegan^40^, bipartite^41^, circlize^42^ and viridis^43^. In R, non-plant sequences were removed from the dataset, as well as the data transformed to relative read abundances (RRAs) per sample. ASVs that were classified as the same plant species were accumulated at the species level. Low abundance taxa that contributed less than 1% to a sample were removed from those samples. The Shannon diversity index was calculated for each sample (pollen and honey) from each bee species. The diversity was tested for significant differences between stingless bee species using the Kruskal-Wallace test, separately for pollen and honey samples. We also performed an NMDS ordination to visualize clustering of samples of pollen and nectar using Bray-Curtis beta-diversity dissimilarities. The ordination represented by proximity of points shows how similar two samples are in terms of composition and abundance of taxa. We tested for differences between species by using a PERMANOVA, separately for honey and pollen samples. We further calculated network indices of the three stingless bee species to account for their overlap and complementarity in the visited plant resources, i.e. the d’ for each bee species and H2’ for the entire network.

## Results

Pollen and honey metabarcoding yielded a total of 5,079,123 quality filtered reads, with mean throughput per sample of 27307.11 reads +/- 1756.635 (SE). Significant reads (more than 1% of reads in any sampling) accounted for 110 ASVs, in 86 genera and 40 plant families; c. 36% of these reads were only matched to generic level or above. In total, 95 out of the 110 ASVs recovered from the samples had been previously recorded in the IBGE Reserve flora^28^; 12 of the 15 absent taxa were exotic cultivated or weedy species. A detailed list of all significant plant species present in pot-pollen and pot-honey samples is available in Table S2. Reads below the threshold value (190 ASVs) still showed a high number of matches to species known to occur in IBGE (86 species, c. 45%) of which 41 were also recorded by us in the RBS floristic inventories.

### How broad is the floral resource exploitation by stingless bees in Cerrado Savanna?

Overall, the interaction network was highly generalized (H2’ = 0.2895575), and consequently also that of the three species within the network (*Melipona rufiventris* d’ = 0.22, *Scaptotrigona postica* d’ = 0.04, and *Tetragonisca angustula* d’ = 0.22) (Figure 2). More than a half of plant species appeared in the samples of at least two of the bee species. In terms of relative plant species abundances as evaluated by combined honey plus pollen samples, bees showed an opportunistic foraging pattern, with most plant species with low abundance and a few highly abundant.

**Figure 2.**
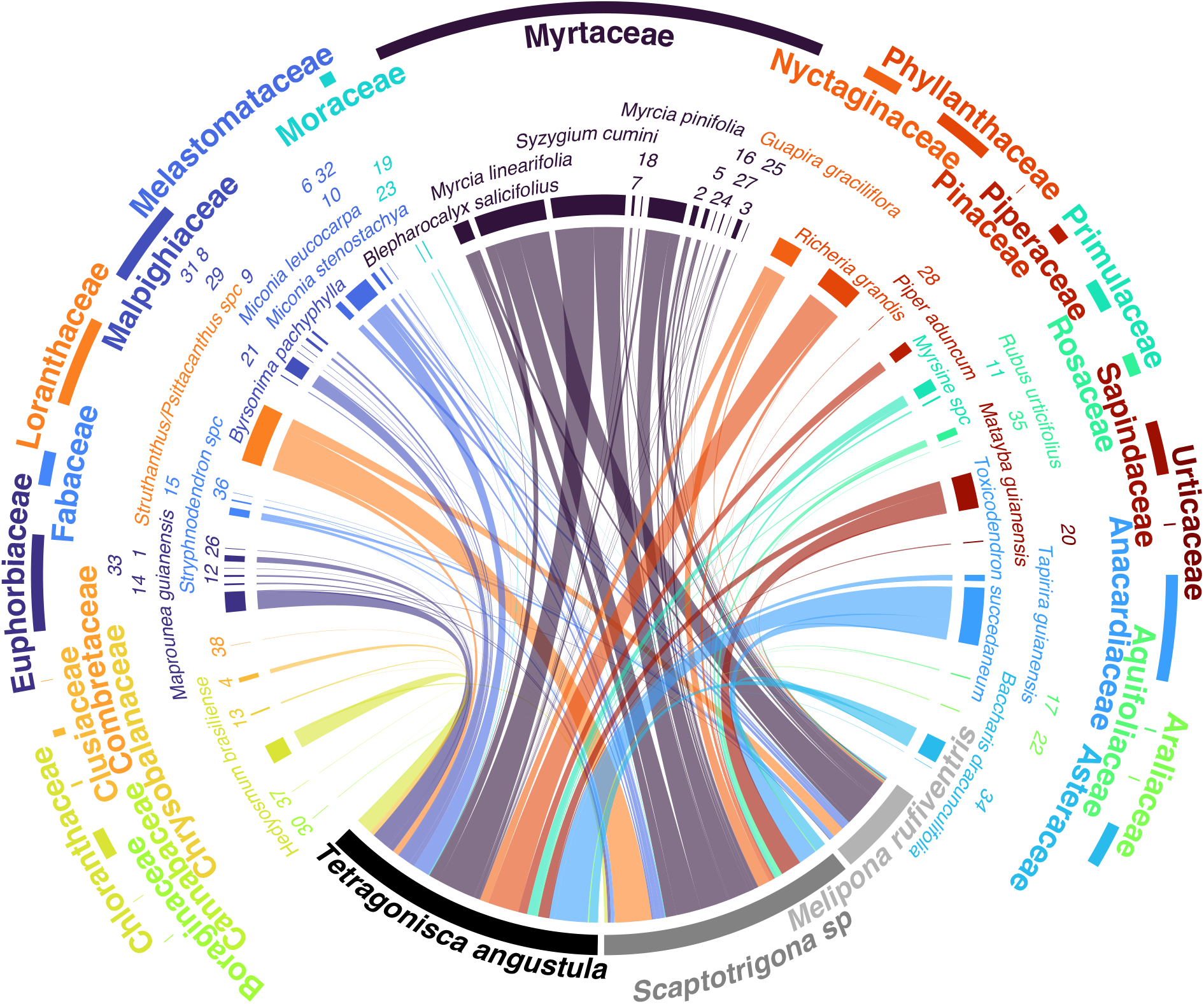
Interaction network of three stingless bee species and the 30 most frequent species in honey and pollen samples (Table S3). Bars connecting bee species and plant species indicate reported interaction (i.e. that plant species was present in the sequencing reads of pollen and/or honey metabarcoding in significant numbers). Some plant species are represented by numbers: 1. *Croton conduplicatus*; 2. *Eucalyptus*; 3. Myrtaceae; 4. *Clusia criuva*; 5. *Myrcia guianensis*; 6. *Miconia hirtella;* 7. *Myrcia splendens*; 8. *Byrsonima basiloba*; 9. *Byrsonima laxiflora*; 10. *Leandra polystachya*; 11. *Myrsine umbellata*; 12. *Acalypha*; 13. *Couepia*; 14. *Mabea fistulifera*; 15. Fabaceae; 16. *Myrcia tomentosa*; 17. *Ilex affinis*; 18. *Eugenia involucrata*; 19. *Moraceae*; 20. *Cecropia pachystachya*; 21. *Byrsonima crassifolia*; 22. *Schefflera macrocarpa*; 23. *Artocarpus heterophyllus*; 24. *Campomanesia pubescens*; 25. *Myrcia pubescens*; 26. *Stillingia*; 27. *Syzygium*; 28. *Pinus*; 29. *Banisteriopsis*; 30. *Borago officinalis*; 31. *Byrsonima viminifolia*; 32. Melastomataceae; 33. *Euphorbia potentilloides*; 34. Asteraceae; 35. *Rosa chinensis*; 36. *Copaifera*; 37. *Trema micranthum*; 38. *Terminalia*.

### Differences among pattern of floral sources exploitation of bee species

The comparison between alpha diversity among samples of different bee species showed that the plant species richness in the pot-honey was higher than in the pot-pollen for all species, but the difference was only significant for *M. rufiventris* (Figure 3). In a comparison among the three bee species, Shannon diversity of plant species in pollen samples was not significantly different between bee species (Kruskal-Wallis rank sum test, chi-squared = 1.4733, df = 2, p-value > 0.05), neither was plant species richness (Kruskal-Wallis rank sum test, chi-squared = 4.5138, df = 2, p-value > 0.05). The same applied for honey samples with Shannon diversity (Kruskal-Wallis rank sum test, chi-squared = 2.6469, df = 2, p-value > 0.05) and species richness (Kruskal-Wallis rank sum test, chi-squared = 4.9389, df = 2, p-value > 0.05).

**Figure 3.**
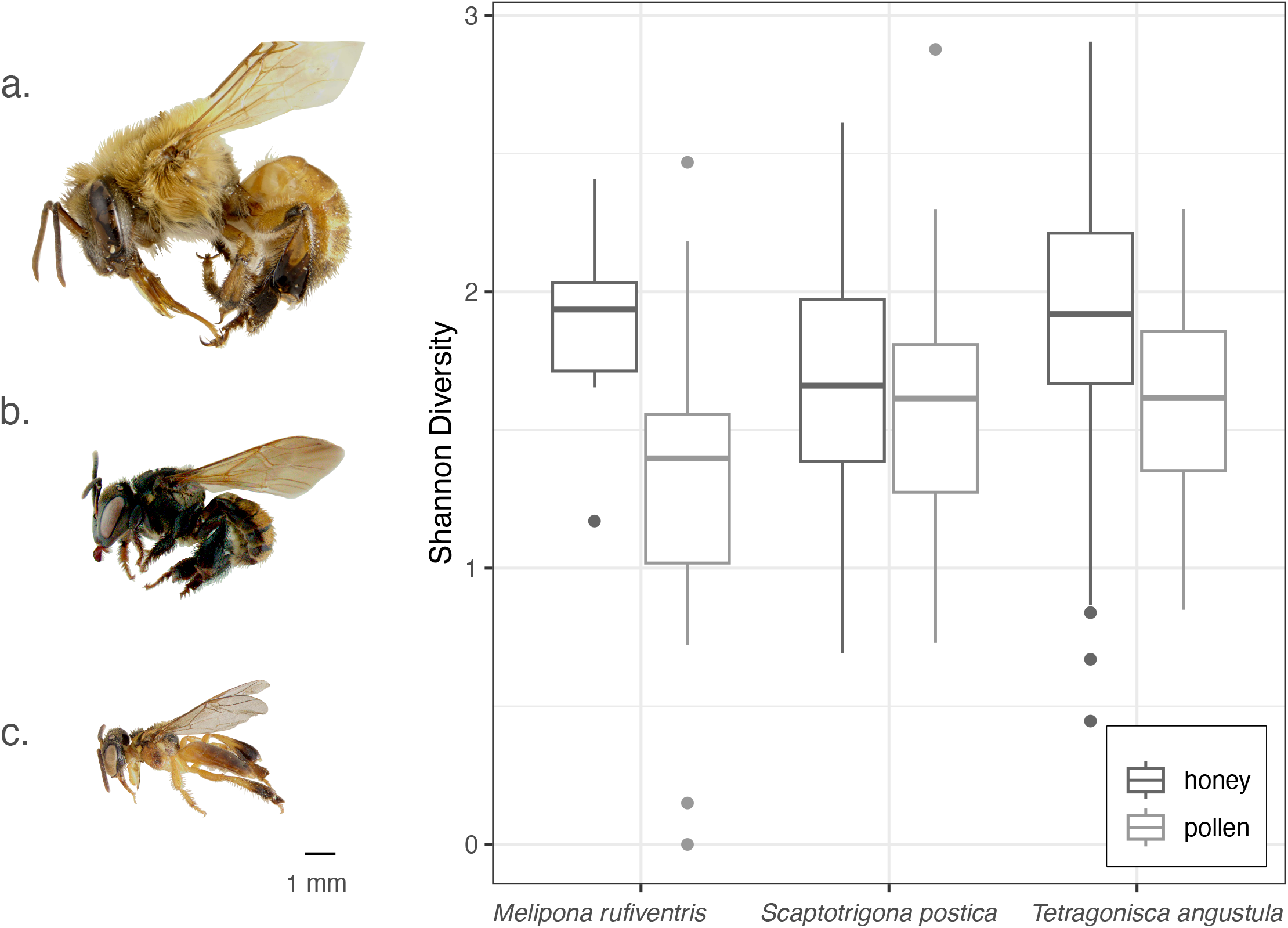
Boxplot of Shannon diversity indexes of plant species found in the honey (dark grey) and pollen (light grey) pots. Boxplots display the median (thick horizontal middle bars), lower (0.25) and upper (0.75) quartile (box limiting thin horizontal bars), minimum and maximum values (vertical lines). Solid dots represent an individual outlier sample. On the left, the three studied bee species in lateral view and in scale to show body size: a. *Melipona rufiventris*, b. *Scaptotrigona postica*, c. *Tetragonisca angustula*.

Although the most frequent plant species are shared among the three stingless bee species, samples from different bee species have several compositional particularities, as shown by the NMDS (Figure 4). The NMDS showed the composition of plants collected differed strongly between bee species, both for pollen (PERMANOVA, df = 2, R2 = 0.12516, F = 7.2246, p < 0.001***) and honey (PERMANOVA, df = 2, R2 = 0.10751, F = 3.8548, p < 0.001***). The NMDS also points to different plant species composition between samples of three species, but in the honey samples little ordination is observed (Fig. 4A). Among pollen samples, on the other hand, we can observe different patterns among the three species, with more overlap between *M. rufiventris* and *S. postica* (Fig. 4B).

**Figure 4.**
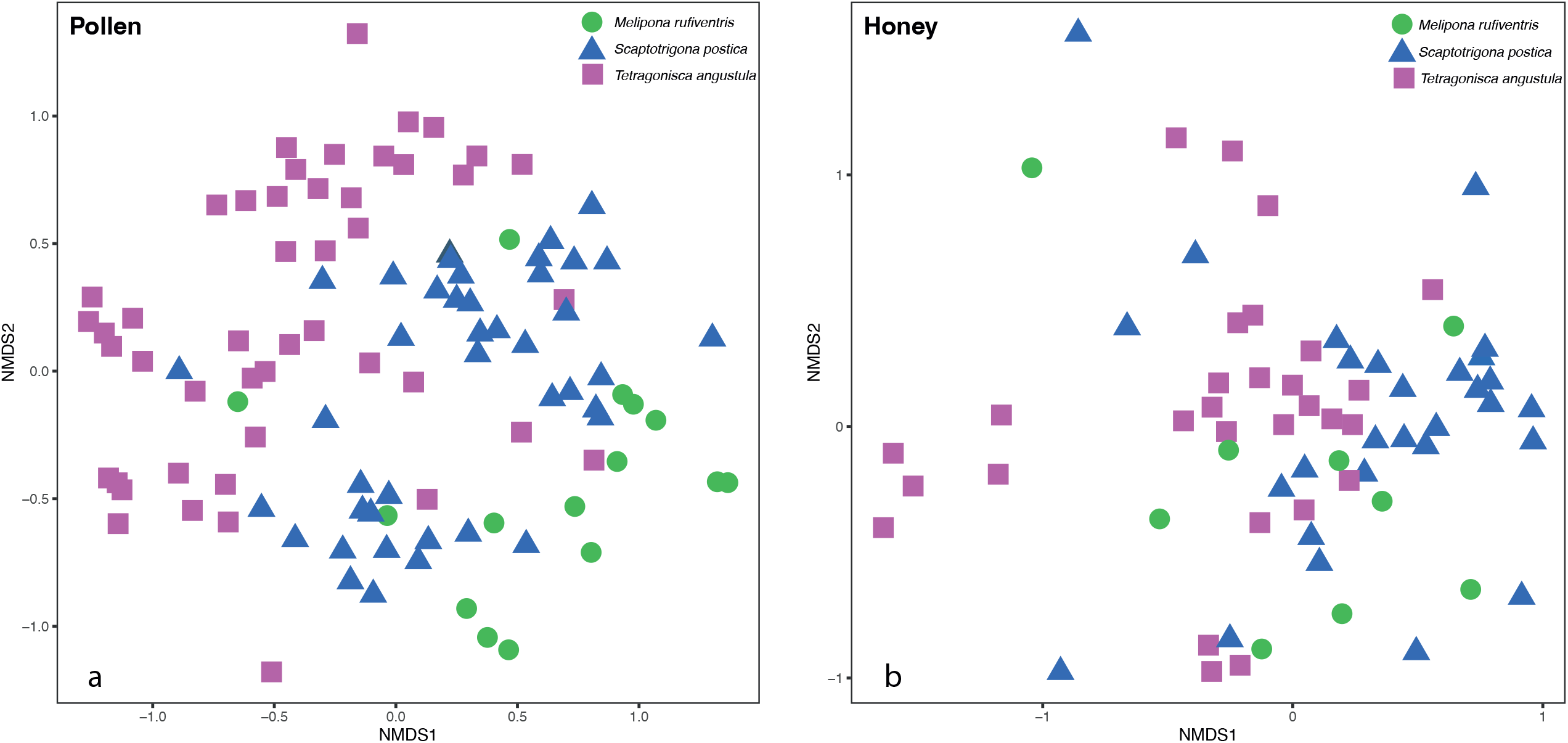
Non-metric multidimensional scaling (NMDS) plots showing plant composition of honey (a) and pot pollen (b) in samples from nests of the three studied bee species: *Melipona rufiventris, Scaptotrigona postica, Tetragonisca angustula.*

### Most frequent plant species and families recovered from pot-pollen and pot-honey samples

The 30 ubiquitously found plant species in pot-honey and pot-pollen samples belong to the following families: Myrtaceae, Loranthaceae, Anacardiaceae, Phyllanthaceae, Sapindaceae, Melastomataceae, Euphorbiaceae, Primulaceae, Nyctaginaceae, Rosaceae, Asteraceae, Malpighiaceae, Cloranthaceae, Piperaceae, Fabaceae, and Clusiaceae (Figure 5, Table S3). Out of 110 ASVs, some plant taxa stand out as most frequent in samples of all the three bee species: Myrtaceae: *Syzygium cumini, Myrcia linearifolia* and *Myrcia pinifolia*; Loranthaceae: *Struthanthus/Psittachanthus*, Anacardiaceae: *Tapirira guianensis*, Phyllanthaceae: *Richeria grandis*, Sapindaceae: *Matayba guianensis*, and Melastomataceae: *Miconia stenostachya*. Most of them offer pollen and nectar, except the pollen-only *Miconia* and the two *Myrcia* species. Thirteen of these ubiquitous species were nectar or oil flowers (i.e., they provide additional resources beyond pollen). Five highly abundant reads were incompletely matched, i.e. could not be identified to species level (*Eucalyptus* sp., Myrtaceae sp., *Myrsine* sp., *Croton* sp, *Struthanthus/Psittachanthus*) but *Croton, Eucalyptus Psittacanthus* and *Struthanthus* are known to produce floral nectar. Pollen-only flowers were found in honey samples of all three species: *Myrsine* sp, *Blepharocalyx salicifolius, Piper aduncum, Miconia leucocarpa* and several *Myrcia* species, thus indicating some kind of mixing nectar and pollen trips, manipulation or spill-over inside the nests. Pollen records include similar diversity numbers of pollen-only flowers and flowers offering nectar and pollen. Only four of out of the 110 ASVs were not recorded in our RBSs: *Bacharis dracunculifolia* and *Myrcia pinifolia*, both native Cerrado species that occur in the IBGE, and exotic *Eucalyptus sp. Toxicodendron succedaneum.*

**Figure 5.**
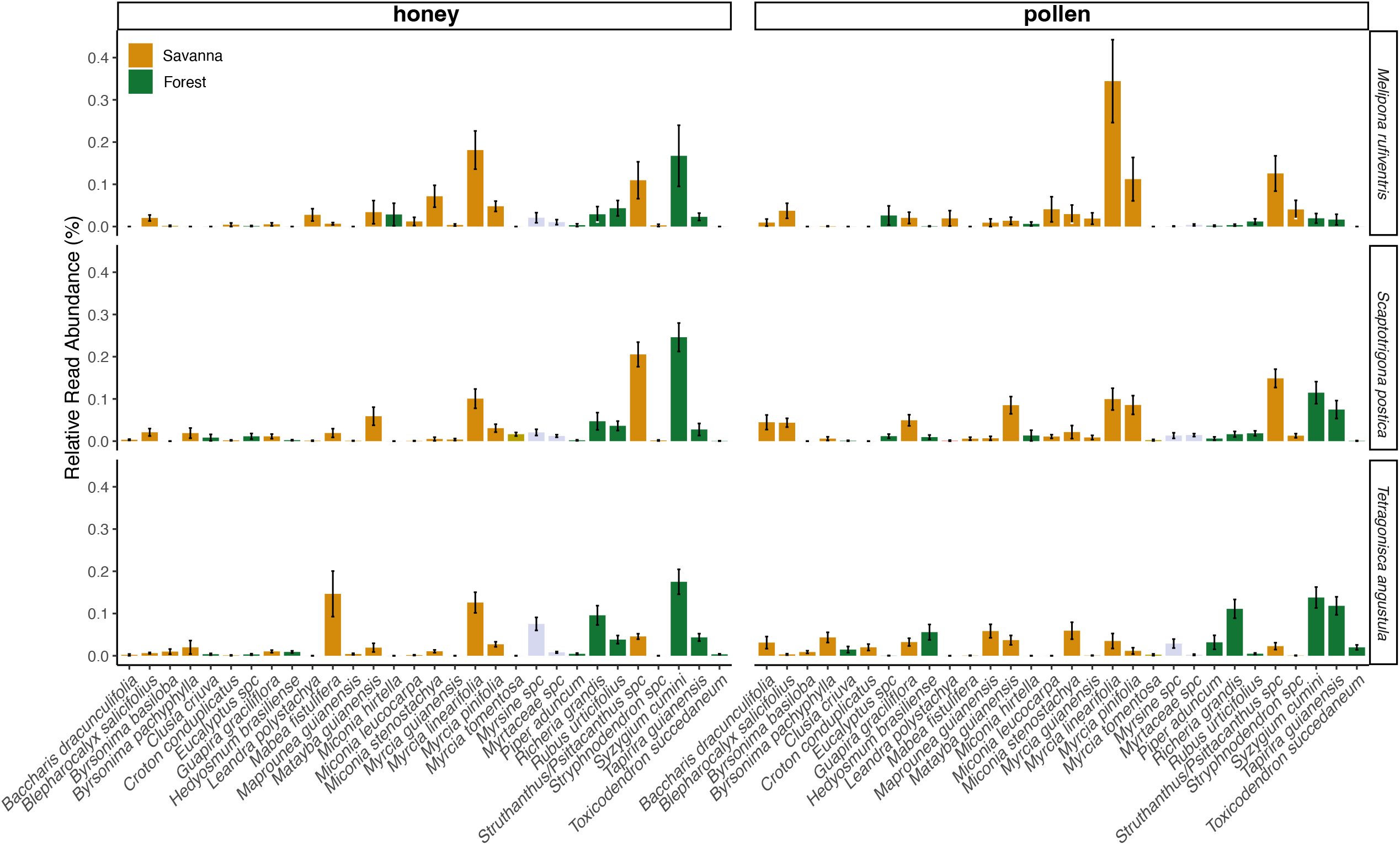
Relative read abundance of the 30 most frequent species found in honey (left half) and pot pollen (right half) samples of nests of three stingless bee species. From top to bottom: *Melipona rufiventris, Scaptotrigona postica, Tetragonisca angustula*. Plant species names are displayed alphabetically. Color in graph bars refers to the habitat of occurrence in Cerrado biome (savanna or forest). Non-identified species were not assigned to any habitat, thus are represented by grey bars.

These 30 most abundant plant species had the following characteristics: all were woody perennials, and most were trees or large shrubs (one climber and one hemiparasite). They could be grouped into two dominant groups according to a combination of the habitat and floral resources. *Group 1* is composed of riverine forest species that offer pollen and nectar, recorded as very common in the Forest RBS surveys: *Syzygium cumini, Tapira guianensis, Richeria grandis, Matayba guianensis, R. urticifolius. Group 2* includes Cerrado shrubs or trees offering only pollen and recorded as common around the nests, in the Cerrado RBS surveys: *Myrcia linearifolia, Blepharocalyx salicifolius, Maprounea guianensis.*

## Discussion

Pollen and honey metabarcoding of three stingless bee species in the genus *Melipona, Scaptotrigona* and *Tetragonisca* revealed a broad generalized set of used floral sources regarding number of species and plant families explored. We recovered 110 plant species in pot-honey and pot-pollen retrieved from nests of three stingless bee species. This reveals a broader spectrum of food sources than found by previous surveys on neotropical stingless bees that relied on non-DNA based methods such as field observations, field collections, and palynological studies. For instance, non-DNA based studies in another hyper diverse area in the Neotropics, the Amazon, revealed from 80 to 122 pollen types in nests and pollen loads of 10-15 species of stingless bees^44^. Other similar studies in species-rich areas of the Neotropics show comparatively lower numbers^22^. While these studies recorded a maximum of five to eight plant species per bee species, we found a mean of 46.3 plant species per bee species in honey samples and 53.67 in pollen samples. The interaction network and high number of species found in honey and pollen of the three analyzed stingless bee species point to a generalist foraging behavior, known to be common in eusocial bees and in stingless bees in particular^17,20^. It also points to probable scouting investigative trips, followed by heavy recruitment and opportunistic behavior when a high-quality resource is located, with most plant species with low abundance and a few highly abundant. Note that our results may still be an underestimation, since samples were collected during only 6 months, i.e., did not include all seasons.

The power of pollen DNA metabarcoding in revealing broad food sources for stingless bees had only been demonstrated before in Southeast Asia and Australia. In Sumatra, a study of *Tetragonula laeviceps* using pollen metabarcoding coupled with light microscopy revealed 99 plant species^45^. Similarly, a study with *Tetragonula carbonaria* in Queensland retrieved 302 plant species in pollen samples across seven sites at different seasons of the year over a two-year period^13^. These are promising results, especially when considering expanding this technique to tropical and subtropical forests of the Neotropics. Studies of pollination and floral biology in these habitats is often very difficult because the plants are scattered, flowers are difficult to reach, and often in the upper canopy. Therefore, direct observations of bees on flowers in tropical and subtropical forests are rare^20^, and records of stingless bee – flower interaction in these environments became almost restricted to pollen loads or pot pollen analyses^22^. Although their utility is undeniable^46^, morphological identification of pollen may become obsolete for pollination biology studies when compared with the efficiency of DNA metabarcoding to identify different plant species in extremely rich floras.

Pollen analyses via DNA metabarcoding also have the advantage of revealing unexpected food sources used by bees that would perhaps be unnoticed in studies using other methodologies. For instance, our analyses revealed that DNA from 13 wind-pollinated plant species were found among the 50 most abundant species in the sample of the three species, including monocots (Poaceae, Cyperaceae), eudicots (Euphorbiaceae: *Acalypha*, Amaranthaceae: *Amaranthus*, Urticaceae: *Cecropia*, Cannabaceae: *Trema*), and a conifer genus, the introduced *Pinus* (Table S2). The presence of non-melitophyllous angiosperms and gymnosperms is relatively common in melissopalynological studies: Cyperaceae, Poaceae, Taxaceae and Pinaceae^22,45,47^. Despite previous studies demonstrating that pollen from anemophilous species might be a contamination in melissopalynological samples ^48^, bees are regularly reported visiting such taxa^49,50^. Our results confirm active collection of pollen from anemophilous species, since their abundance in our analyzed samples is relatively high. One of the most abundant plant species in the pollen analysis was *Hedyosmum brasiliense* (Chloranthaceae), widely cited in the literature as wind-pollinated^51^. This species was not only recorded in the pollen samples of all three species of bees, but was amongst the 10 most abundant records for *Tetragonisca angustula* in our results. These results reinforce the theory that anemophilous plants, which account for 10% of angiosperms and most gymnosperms, produce enough pollen^52^ to be attractive to social bees, under certain conditions of colony size and food demands. However, the role of bees and other insects as true pollinators of anemophilous plants remains unresolved, in spite of the importance of wind-pollinated crops^53^ and of the several records showing that anemophilous plant pollen is important for several bee species (see references above).

A surprising and novel observation is the significant amount of Marchanthyophyte DNA from the liverwort *Dumortiera hirsuta* found in pot pollen from the three studied stingless bee species (Table S2). Future research would need to seek evidence if the DNA results from the collection of spores or perhaps some chemical compounds from liverworts by stingless bees. Bees collecting spores from fungi and plants is not a novelty, as there is evidence of active collecting^54^ as well as records of spores in samples of pollen and honey^55^. In lieu of pollen, spores supposedly have nutritional benefits^56^. Stingless bees might also visit liverworts to collect lipidic compounds, e.g. terpenoids used in communication among individuals^57^ commonly occurring in liverworts^58^.

The high degree of overlap between plant profiles found in the honey of the three bee species suggests that bees may be competing for the same nectar resources. Pollen plant profiles on the other hand showed far less overlap between species, corroborating evidence that pollen exploitation and digestion requires a high degree of specialization^59^, even in generalist bees^60^, which is often facilitated by each species’ microbiome^61^. Although some plant species appeared in the samples of all three bee species, *Scaptotrigona postica* and *M. rufiventris* shared more species while *T. angustula* differed from both. Considering body size vs. flower matching, the smallest species, *T. angustula*, visits the highest number of species of the three, potentially due to solitary foraging behavior, in which females forage alone without recruiting other workers.

*Melipona* species present a unique foraging pattern among stingless bees, not only because they are amongst the largest stingless bees (up to 15 mm, Michener 2007), but because they show clear preferences towards some groups of plants^22,62^. *Melipona* are also the only stingless bees capable of buzzing to harvest pollen^63^, but pollen-flowers that require buzz-pollination for pollen harvesting were not abundant in the samples, even though species with poricidal anthers were observed flowering around the nests during the months of collection (e.g., *Miconia ferruginata* DC, *Pleroma stenocarpum* (Schrank & Mart. ex DC.) Triana, *Solanum falciforme* Farruggia).

Our botanical surveys also reinforced the patterns of floral exploitation among the three species, such as the apparent preference for trees with mass flowering by stingless bees, even though their exploitation demands a long flight range. Some stingless bees’ sophisticated communication abilities allow a massive recruitment of foragers when mass blooming plants are available^19^. In the case of *Tetragonisca angustula*, which is considered a solitary forager, the range of pollen sources is wider and seems less biased towards mass blooming plants. In the Atlantic rainforest, another hyper diverse neotropical ecosystem,^20^ observed that stingless bees have a preference for upper canopy stratum with small hermaphroditic or monoecious whitish flowers and abundant resources (pollen and/or nectar). Importantly, most of their preferred trees flower in mass, i.e produce a large number of flowers over a short period of time^20^. In the Cerrado savannas, where the nests were, we observed the typical high frequency of shrubs and herbaceous species in stingless bees pollen (ca. 38% of samples), which reflects the savanna physiognomy where herbs and shrubs are predominant^64^. However, despite the high availability of flowers in the savanna surrounding their nests, they still flew up to riverine forests at least 630 m far from the nests to collect resources where mass-flowering species were more common.

Flight distance in bees is usually related to body size (larger bees tend to have wider flight ranges)^65^ and social behavior (social bees have a larger foraging distance than solitary bees due to the potential communication and recruitment between individuals)^66^. Given that the closest riverine forest is located at a distance of 630 m to the nests, and that species from this habitat were among the most abundant in the samples, this suggests that all three stingless bee species will forage and probably recruit at least 630 m from their nests, supporting the hypothesis of long-distance foraging when attractive rewards are available^20^. This distance is well within the known flight range of *Melipona* whose typical flight distance is about 2 km, but can be extended up to 10 km^18^, but it is more surprising for *Scaptotrigona* and *Tetragonisca* whose reported maximum flight distances are 1.7 to 0.6 km, respectively^18^.

These estimates of minimum foraging distance of 630m are considered trustworthy based on the high frequency of pollen from species occurring only in riverine forests (Group 1), e.g. *Syzygium cumini*, an introduced species that only occurs in a small portion of the nearest riverine forest to the nests. Other highly abundant species in our samples are common in the Distrito Federal riverine forests (*Clusia cruiva, Hedyosmum brasiliense, Miconia hirtella, Piper aduncum, Richeria grandis*)^67–69^ and were only found in our surveys of the riverine forests (Table 1).

Some plant families stand out as the most important floral sources for the three stingless bee species, i.e. have one or more species amongst the 30 most frequent ASVs. Amongst them, Myrtaceae, Anacardiaceae, Sapindaceae, Melastomataceae, Euphorbiaceae, and Asteraceae are well-known as common resources for stingless bees globally^17^, while Loranthaceae and Malpighiaceae are frequent in other studies^62^. Phyllanthaceae, Primulaceae, Chloranthaceae and Piperaceae, however, have been only rarely reported^22^. Asteraceae, Myrtaceae, and Melastomataceae are amongst the most speciose plant families in the IBGE reserve, representing at least 300 species with different life forms (from herbs to trees) in the flora^28^, but it is surprising that other diverse plant families in the IBGE area, i.e. Fabaceae, Lamiaceae and Orchidaceae, which also represent close to 300 species combined^28^, are less conspicuous or totally absent from our most frequent 30 taxa. This means that, although the important floral sources for stingless bees partially overlap with the most common plants in the area, indicating that abundant sources are preferred, this is not always the case. This could simply mean that species within these families were not flowering at the time of sampling, but it is worth noting that Lamiaceae, papilionoid legumes and orchids share complex floral morphologies that are different from those of the families recorded as most abundant in our samples These three families tend to present flowers with bilateral symmetry, specialized petals and androecia, and deep, hidden resources that often forces floral visitors to approach and handle the flowers in a specific way^70^. Our results confirm the hypothesis raised by^20^ that stingless bees may be specialized in exploiting small, open resource “bowl-type” flowers^52^, with exposed stamens and nectar, that are produced in large numbers. They may also favour plant species with a “big bang” flowering phenology i.e., that that undergo mass blooming for short periods. Floral morphology, floral chemistry and phenology of plants exploited by stingless bees deserve further investigation. Investigations of plant resources exploited by stingless bees using metabarcoding over a longer time periods, in other types of vegetation, and of other bee species, would also be desirable to consolidate our knowledge of stingless bee ecology in the Neotropics.

## Acknowledgements

We would like to thank Instituto Serapilheira for the grant conceived (Chamada Publica N° 2 - 2018). We thank the IBGE reserve for allowing and supporting our field work collecting data; Antonio Leite from IBRAMEL stingless beekeepers association for kindly donating the nests that were used in the experiments. AJCA acknowledged FAPDF for the financial support (00193-00001229/2021-48). ACM thanks the CNPq for the postdoctoral fellowship (159694/2018-3) and CEBP for the CNPQ PQ2 fellowship. KN & MMC thanks their undergraduate scholarships (ProIC UnB). FAC acknowledge the CAPES/PRINT program (Edital nº 41/2017 88887.716844/2022-00) for allowing the visit of Alexander Keller to UFMG.

## Data Availability

The data that support the findings of this study is available at public repositories: molecular sequence data is available at NCBI (Bioproject 976708); plant species list with voucher information is available at Species Link by searching on the collector name “Projeto Barcode Cerrado”.

## Supplementary information

**Table S1.** Pollen and honey sampling collected from bee nests of the three stingless bee species: *Melipona rufiventris* (M), *Scaptotrigona postica* (S) and *Tetragonisca angustula* (T).

**Table S2.** Amplicon sequences varieties (ASVs) with significant number of reads and their taxon matches. The IBGE column records presence/absence of taxa of any level in the IBGE flora (IBGE 2011). The RBS column records if/where species were recorded in the floristic survey (distances given from nests): G=garden (650m); N=nest plot (50m); I=inner pentagon plots (700m); O=outer pentagon plots (1500m); F1=near forest (630m); F2= distant forest (2070); ? = automatically attributed to all reads not matched to species. The occurrence in honey or pollen is indicated by the bee species acronym in the relevant column: MR, *Melipona rufiventris*; SP*, Scaptotrigona postica* and TA, *Tetragonisca angustula*. Floral rewards to pollinators (pollen, nectar or oil) is presented as well as if the species is traditionally considered wind-pollinated. We assume all non-wind-pollinated are animal pollinated plants.

## References

1. Klein, A. M. et al. Importance of pollinators in changing landscapes for world crops. Proceedings. Biological sciences / The Royal Society274, 303–13 (2007).

2. Gallai, N., Salles, J., Settele, J. & Vaissiere, B. Economic valuation of the vulnerability of world agriculture confronted with pollinator decline. Ecological Economics68, 810–821 (2009).

3. Michener, C. D. The bees of the world. (The John Hopkins University Press, 2007).

4. Ollerton, J., Winfree, R. & Tarrant, S. How many flowering plants are pollinated by animals? Oikos120, 321–326 (2011).

5. Biesmeijer, J. C. et al. Parallel declines in pollinators and insect-pollinated plants in Britain and the Netherlands. Science(1979) 313, 351–354 (2006).

6. Morales, C. L. et al. Does climate change influence the current and future projected distribution of an endangered species? The case of the southernmost bumblebee in the world. J Insect Conserv26, 257–269 (2022).

7. Parreño, M. A. et al. Critical links between biodiversity and health in wild bee conservation. Trends Ecol Evol37, 309–321 (2022).

8. Keller, A. et al. Evaluating multiplexed next-generation sequencing as a method in palynology for mixed pollen samples. Plant Biol17, 558–566 (2015).

9. Sickel, W. et al. Increased efficiency in identifying mixed pollen samples by meta-barcoding with a dual-indexing approach. BMC Ecol15, 1–9 (2015).

10. Baksay, S. et al. Experimental quantification of pollen with DNA metabarcoding using ITS1 and trnL. Sci Rep10, (2020).

11. Khansaritoreh, E. et al. Employing DNA metabarcoding to determine the geographical origin of honey. Heliyon6, (2020).

12. Elliott, B. et al. Pollen diets and niche overlap of honey bees and native bees in protected areas. Basic Appl Ecol50, 169–180 (2021).

13. Wilson, R. S. et al. Landscape simplification modifies trap-nesting bee and wasp communities in the subtropics. Insects11, 1–15 (2020).

14. Camargo, J. M. F., Pedro, S. R. M. & Melo, G. A. R. Meliponini Lepeletier, 1836. in Catalogue of Bees(Hymenoptera, Apoidea) in the Neotropical Region - online version. (2013).

15. Janousek, W. M. et al. Recent and future declines of a historically widespread pollinator linked to climate, land cover, and pesticides. Proc Natl Acad Sci U S A120, (2023).

16. Slaa, E. J., Chaves, L. A. S., Malagodi-Braga, K. & Hofstede, F. E. Stingless bees in applied pollination: practice and perspectives. Apidologie37, 293–315 (2006).

17. Bueno, F. G. B. et al. Stingless bee floral visitation in the global tropics and subtropics. Glob Ecol Conserv43, e02454 (2023).

18. Nunes-Silva, P. et al. Radiofrequency identification (RFID) reveals long-distance flight and homing abilities of the stingless bee Melipona fasciculata. Apidologie51, 240–253 (2020).

19. Biesmeijer, J. C. & Slaa, E. J. Information flow and organization of stingless bee foraging. Apidologie35, 143–157 (2004).

20. Ramalho, M. Stingless bees and mass flowering trees in the canopy of Atlantic Forest: a tight relationship. Acta Bot Brasilica18, 37–47 (2004).

21. Bell, K. L. et al. Pollen DNA barcoding□: current applications and future. Genome640, 629–640 (2016).

22. Vit, P., Pedro, S. R. M. & Roubik, D. W. Pot-pollen in stingless bee melittology. Pot-Pollen in Stingless Bee Melittology(Springer International Publishing, 2018). doi:10.1007/978-3-319-61839-5.

23. Myers, N., Mittermeier, R. A., Mittermeier, C. G., da Fonseca, G. A. B. & Kent, J. Biodiversity hotspots for conservation priorities. Nature403, 853 (2000).

24. Flora e Funga do Brasil. Flora e Funga do Brasil. Jardim Botanico do Rio de Janeirohttps://floradobrasil.jbrj.gov.br/.

25. Silva-Souza, K. J. P., Pivato, M. G., Silva, V. C., Haidar, R. F. & Souza, A. F. New patterns of the tree beta diversity and its determinants in the largest savanna and wetland biomes of South America. Plant Divers(2022) doi:10.1016/j.pld.2022.09.006.

26. Silberbauer-Gottsberger, I. & Gottsberger, G. A polinização de plantas do Cerrado. Rev Bras Biol48, 651–663 (1988).

27. Vasconcelos, T. A trait-based approach to determining principles of plant biogeography. Am J Bot110, (2023).

28. Pereira, B. A. S. & Furtado, P. P. Vegetação da bacia do córrego Taquara: coberturas naturais e antrópicas. in Reserva Ecológica do IBGE: Biodiversidade Terrestre vol. 1 89–117 (2011).

29. Rasmussen, C. & Cameron, S. A. Global stingless bee phylogeny supports ancient divergence, vicariance, and long distance dispersal. Biological Journal of the Linnean Society99, 206–232 (2010).

30. Campos, M. G. et al. Standard methods for pollen research. J Apic Res60, 1–109 (2021).

31. Rognes, T., Flouri, T., Nichols, B., Quince, C. & Mahé, F. VSEARCH: a versatile open source tool for metagenomics. PeerJ4, e2584 (2016).

32. Edgar, R. C. & Flyvbjerg, H. Error filtering, pair assembly and error correction for next-generation sequencing reads. Bioinformatics31, 3476–3482 (2015).

33. Edgar, R. C. UCHIME2: improved chimera prediction for amplicon sequencing. bioRxiv074252 (2016) doi:10.1101/074252.

34. Keller, A. et al. BCdatabaser: on-the-fly reference database creation for (meta-) barcoding. Bioinformatics36, 2630–2631 (2020).

35. Edgar, R. C. SINTAX: a simple non-Bayesian taxonomy classifier for 16S and ITS sequences. bioRxiv074161 (2016) doi:10.1101/074161.

36. Ankenbrand, M. J., Keller, A., Wolf, M., Schultz, J. & Förster, F. ITS2 Database V: twice as much. Mol Biol Evol32, 3030–3032 (2015).

37. Marshall, C. A. M., Wieringa, J. J. & Hawthorne, W. D. Bioquality hotspots in the tropical African flora. Current Biology26, 3214–3219 (2016).

38. R Core Team. R: a language and environment for statistical computing. Preprint at https://www.r-project.org/ (2021).

39. McMurdie, P. J. & Holmes, S. phyloseq: An R Package for reproducible interactive analysis and graphics of microbiome census data. PLoS One8, e61217.(2013).

40. Oksanen, J., Kindt, R., Legendre, P., O’Hara, B. & Stevens, H. The vegan package. Community ecology package10, 631–637 (2007).

41. Dormann, C. F., Gruber, B. & Fründ, J. Introducing the bipartite Package: Analysing Ecological Networks. R News8, 8–11 (2008).

42. Gu, Z., Gu, L., Eils, R., Schlesner, M. & Brors, B. circlizeimplements and enhances circular visualization in R. Bioinformatics30, 2811–2812 (2014).

43. Garnier, S. et al. viridis-Colorblind-Friendly Color Maps for R. Preprint at (2021).

44. Absy, M. L., Rech, A. R. & Ferreira, M. G. Pollen Collected by Stingless Bees: A Contribution to Understanding Amazonian Biodiversity. in Pot-Pollen in Stingless Bee Melittology29–46 (Springer International Publishing, 2018). doi:10.1007/978-3-319-61839-5_3.

45. Moura, C. C. M. et al. Biomonitoring via DNA metabarcoding and light microscopy of bee pollen in rainforest transformation landscapes of Sumatra. BMC Ecol Evol22, 1–15 (2022).

46. Roubik, D. & Patiño, J. E. M. The stingless honey bees (Apidae, Apinae: Meliponini) in Panama and pollination ecology from pollen analysis. in Pot-Pollen in Stingless Bee Melittology(eds. Vit, P., Pedro, S. R. M. & Roubik, D. W.) 47–66 (Springer, 2018).

47. Elliott, B. et al. Pollen diets and niche overlap of honey bees and native bees in protected areas. Basic Appl Ecol50, 169–180 (2021).

48. Pound, M. J. et al. Determining if honey bees (Apis mellifera) collect pollen from anemophilous plants in the UK. Palynology(2022) doi:10.1080/01916122.2022.2154867.

49. Malerbo-Souza, D. T., Da Silva, T. G., De Andrade, M. O., De Farias, L. R. & Medeiros, N. M. G. Factors affecting the foraging behavior of bees in different maize hybrids. Revista Brasileira de Ciencias Agrarias13, (2018).

50. Costa, A. C. G., Albuquerque, I. S., Thomas, W. W. & Machado, I. C. Influence of environmental variation on the pollination of the ambophilous sedge Rhynchospora ciliata(Cyperaceae). Plant Ecol219, 241–250 (2018).

51. Gottsberger, G. Generalist and specialist pollination in basal angiosperms (ANITA grade, basal monocots, magnoliids, Chloranthaceae and Ceratophyllaceae): what we know now. Plant Divers Evol131, 263–362 (2015).

52. Faegri, K. & van der Pijl, L. Principles of pollination ecology. (Pergamon Press, 1979).

53. Saunders, M. E. Insect pollinators collect pollen from wind-pollinated plants: implications for pollination ecology and sustainable agriculture. Insect Conserv Divers11, 13–31 (2018).

54. Oliveira, M. L. & Morato, E. F. Stingless bees (Hymenoptera, Meliponini) feeding on stinkhorn spores (Fungi, Phallales): robbery or dispersal? Rev Bras Zool 17, 881–884 (2000).

55. Barth, O. M., Freitas, A. da S. de & Rio Branco, C. dos S. Pollen collected by stingless bees in a reforested urban area of Rio de Janeiro city. Bee World98, 23–26 (2021).

56. Parish, J. B., Scott, E. S. & Hogendoorn, K. Nutritional benefit of fungal spores for honey bee workers. Sci Rep10, (2020).

57. Leonhardt, S. D. Chemical ecology of stingless bees. J Chem Ecol43, 385–402 (2017).

58. Asakawa, Y. Highlights in phytochemistry of hepaticae-biologically active terpenoids and aromatic compounds. Pure & Appl. Chem66, 2193–2196 (1994).

59. Sedivy, C., Müller, A. & Dorn, S. Closely related pollen generalist bees differ in their ability to develop on the same pollen diet: Evidence for physiological adaptations to digest pollen. Funct Ecol25, 718–725 (2011).

60. Bryś, M. S., Skowronek, P. & Strachecka, A. Pollen diet—properties and impact on a bee colony. Insects12, (2021).

61. Keller, A. et al. (More than) Hitchhikers through the network: The shared microbiome of bees and flowers. Curr Opin Insect Sci44, 8–15 (2021).

62. Ramalho, M., Kleinert-Giovannini, A. & Imperatriz-Fonseca, V. L. Important bee plants for stingless bees (Melipona and Trigonini) and africanized honeybees (Apis mellifera) in neotropical habitats: a review. Apidologie21, 469–488 (1990).

63. Nunes-Silva, P., Hrncir, M., Da Silva, C. I., Roldão, Y. S. & Imperatriz-Fonseca, V. L. Stingless bees, Melipona fasciculata, as efficient pollinators of eggplant (Solanum melongena) in greenhouses. Apidologie44, 537–546 (2013).

64. Klink, C. A., Sato, M. N., Cordeiro, G. G. & Ramos, M. I. M. The role of vegetation on the dynamics of water and fire in the cerrado ecosystems: Implications for management and conservation. Plants vol. 9 1–27 Preprint at https://doi.org/10.3390/plants9121803 (2020).

65. Greenleaf, S. S., Williams, N. M., Winfree, R. & Kremen, C. Bee foraging ranges and their relationship to body size. Oecologia153, 589–96 (2007).

66. Grüter, C. & Hayes, L. Sociality is a key driver of foraging ranges in bees. Current Biology32, 5390-5397.e3 (2022).

67. Darosci, A. A. B., Takahashi, F. S. C., Proença, C. E. B., Soares-Silva, L. H. & Munhoz, C. B. R. Does spatial and seasonal variability in fleshy-fruited trees affect fruit availability? A case study in gallery forests of Central Brazil. Acta Bot Brasilica35, 456–465 (2021).

68. Mendonça, R. C. et al./person-group>. Flora vascular do Cerrado. in Cerrado: ambiente e flora(eds. Sano, S. M. & Almeida, S. P.) 289–556 (Embrapa-Cerrados, 1998).

69. Ratter, J. A., Bridgewater, S. & Ribeiro, F. Analysis of the floristic composition of the Brazilian cerrado vegetation III: comparison of the woody vegetation of 376 areas. Edinb J Bot60, 57–109 (2003).

70. Willmer, P. Pollination and floral ecology. in Pollination and floral ecology (Princeton University Press, 2011).

